# Coronary Artery Disease Risk Gene *PRDM16* is Preferentially Expressed in Vascular Smooth Muscle Cells and a Potential Novel Regulator of Smooth Muscle Homeostasis

**DOI:** 10.1101/2023.04.03.535461

**Authors:** Kunzhe Dong, Xiangqin He, Guoqing Hu, Yali Yao, Jiliang Zhou

## Abstract

**Objective:** Vascular smooth muscle cells (VSMCs) are the primary contractile component of blood vessels and can undergo phenotypic switching from a contractile to a synthetic phenotype in vascular diseases such as coronary artery disease (CAD). This process leads to decreased expression of SMC lineage genes and increased proliferative, migratory and secretory abilities that drive disease progression. Super-enhancers (SE) and occupied transcription factors are believed to drive expression of genes that maintain cell identify and homeostasis. The goal of this study is to identify novel regulator of VSMC homeostasis by screening for SE-regulated transcription factors in arterial tissues.

**Approach and Results:** We characterized human artery SEs by analyzing the enhancer histone mark H3K27ac ChIP-seq data of multiple arterial tissues. We unexpectedly discovered the transcription factor PRDM16, a GWAS identified CAD risk gene with previously well-documented roles in brown adipocytes but with an unknown function in vascular disease progression, is enriched with artery-specific SEs. Further analysis of public bulk RNA-seq and scRNA-seq datasets, as well as qRT-PCR and Western blotting analysis, demonstrated that PRDM16 is preferentially expressed in arterial tissues and in contractile VSMCs but not in visceral SMCs, and down-regulated in phenotypically modulated VSMCs. To explore the function of *Prdm16* in vivo, we generated *Prdm16* SMC-specific knockout mice and performed histological and bulk RNA-Seq analysis of aortic tissues. SMC-deficiency of *Prdm16* does not affect the aortic morphology but significantly alters expression of many CAD risk genes and genes involved in VSMC phenotypic modulation. Specifically, *Prdm16* negatively regulates the expression of *Tgfb2* that encodes for an upstream ligand of TGF-β signaling pathway, potentially through binding to the promoter region of *Tgfb2*. These transcriptomic changes likely disrupt VSMC homeostasis and predispose VSMCs to a disease state.

**Conclusions:** Our results suggest that the CAD risk gene *PRDM16* is preferentially expressed in VSMCs and is a novel regulator of VSMC homeostasis. Future studies are warranted to investigate its role in VSMCs under pathological conditions such as atherosclerosis.

## INTRODUCTION

Smooth muscle cells (SMCs) are the major contractile components in hollow organs such as blood vessels and gastrointestinal (GI) tissues. Under normal conditions, SMCs are highly differentiated and contractile, characterized by expressing a unique repertoire of contractile genes ^1^. However, under pathological conditions such as coronary artery disease (CAD), SMCs switch their phenotype from a contractile to a synthetic state, as characterized by decreased expression of SMC-specific lineage genes, increased expression of genes associated with proliferation, migration and secretion ^1^. This process is known as SMC phenotypic modulation and plays a major role in driving vascular disease progression ^2^. However, the mechanism governing SMC phenotypic modulation is largely unknown. Identifying novel players that maintains SMC identity and homeostasis is an essential step towards understanding the etiology of vascular diseases and developing better clinical approaches.

Super-enhancers (SEs) and occupied transcription factors (TFs) are predominant determinants of cell identify and homeostasis by precisely controlling the spatiotemporal expression of cell lineage-specific genes ^3-5^. Recent studies have shown that expression of many critical cardiac TFs, such as TBX3 ^6^, GATA4 ^7^, MESP1/2 ^8^ and HAND2 ^9^, is driven by cardiac SEs. Moreover, sequence mutations occurring within the enhancer regions can alter the expression of cardiac genes and cause cardiac dysfunction ^10-15^. These studies implicate the potential of identifying novel regulators that maintain SMC identity and hemostasis by screening for vascular SMC (VSMC)-specific SE-regulated TFs.

PRDM16 is a member of the PR domain-containing (PRDM) family and encodes a zinc-finger TF ^16^. Mutations of *PRDM16* gene have been associated with various cardiovascular diseases including CAD ^17^, blood pressure ^18,19^, stroke ^20^ and dilated cardiomyopathy ^21,22^. However, the underlying mechanisms for its involvement in disease progression is poorly understood, such as its in vivo expression pattern and biological function ^23^. PRDM16 is well known as a master mediator of brown fat differentiation and identify ^24-26^. Recently, PRDM16 is shown to play a critical role in maintaining myocardial cardiomyocyte identity ^27^ and regulating arterial flow recovery by maintaining endothelial function ^28^. However, its role in VSMCs is almost entirely unknown.

Through an unbiased genome-wide screening of multiple public epigenetic and transcriptomic datasets, here we unexpectedly discovered that *PRDM16* is enriched with artery-specific SEs and preferentially expressed in arterial tissues and VSMCs. We further explored the PRDM16 function in VSMCs at baseline by generating *Prdm16* inducible SMC-specific KO (iSM KO) mice and performing histological and whole transcriptome analysis of aortic tissues. Our results suggest that loss of *Prdm16* in SMCs does not affect aortic morphology but significantly alters expression of genes involved in cardiovascular disease progression, response to external stimuli, cell proliferation and migration, as well as TGF-β (transforming growth factor beta) signaling pathway upstream ligand *Tgfb2* gene. We propose these transcriptomic changes resulted from *Prdm16* deficiency disrupt VSMC homeostasis and predispose VSMCs to a disease sate.

## MATERIALS AND METHODS

The authors declare that all supporting data are included in the article and its online Supplemental Material. All animal procedures were performed in accordance with the National Institutes of Health Guide for the Care and Use of Laboratory Animals and approved by the Institutional Animal Care and Use Committee of Augusta University.

## RESULTS

### The CAD risk gene *PRDM16* is preferentially expressed in arterial tissues and VSMCs

To establish the epigenetic regulatory atlas of human blood vessels, we de novo analyzed the enhancer data of 9 human arterial tissues defined by H3K27ac ChIP-seq from ENCODE database (**Online Table S1**). We characterized human artery SEs with the established ROSA algorithm for SE analysis ^3^ and integrated the highly reliable SEs (present in more than 5 samples) with human TF database to screen for TFs that are regulated by SEs in human arteries (**Figure 1A**). We prioritized the obtained 1,071 arterial SEs based on a combination of average ranking and intensity value (**Figure 1B**). This analysis revealed 126 SE-regulated TFs in human arterial tissues. Among them, PRDM16, which is a GWAS identified CAD risk gene with unknown function in the progression of vascular disease ^17,23^, stood out as having the smallest average ranking number and strong intensity across the 9 samples (**Figure 1B**).

**Figure 1.**
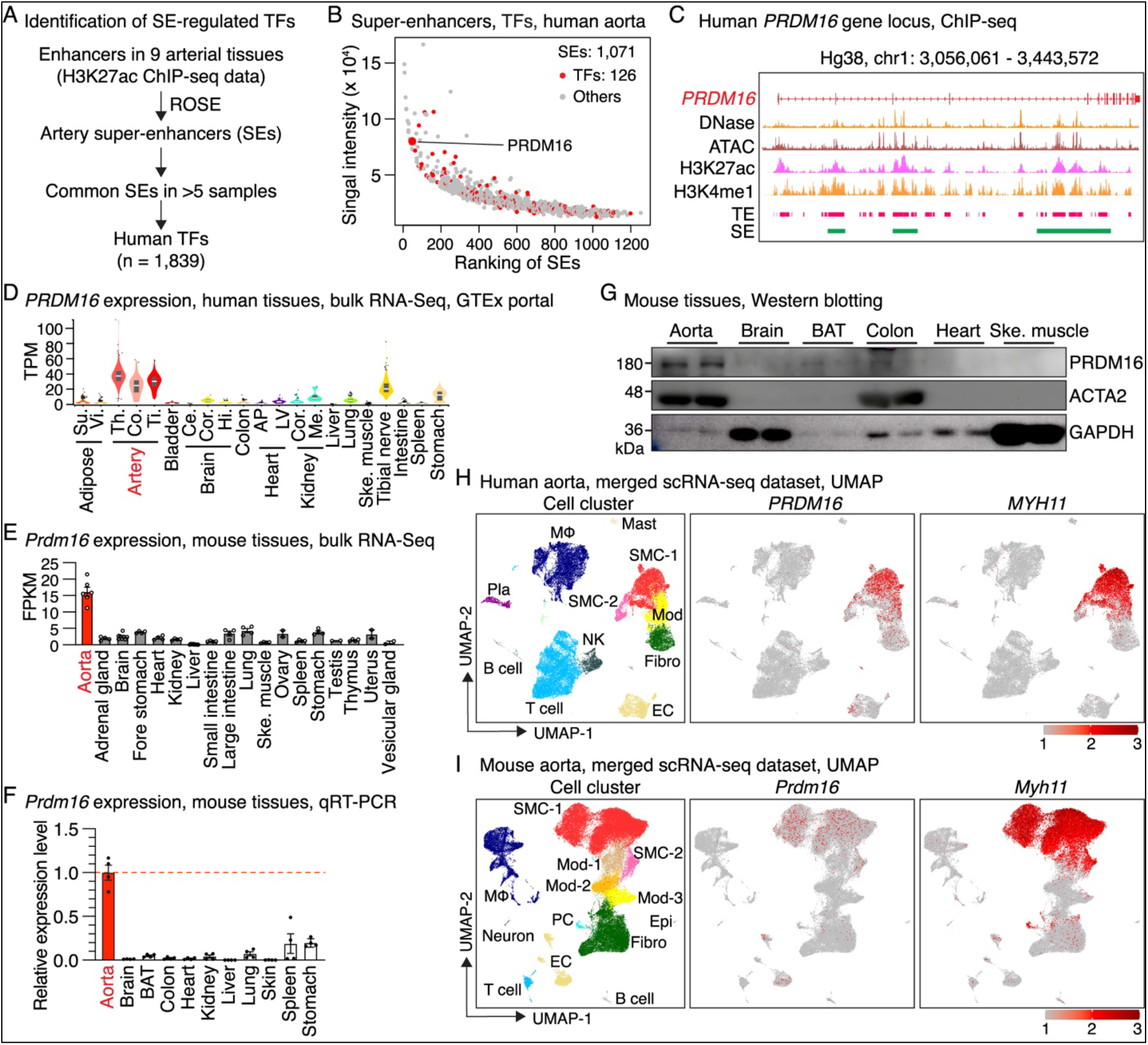
PRDM16 is preferentially expressed in arterial tissues and VSMCs. **(A)** Workflow for identifying super-enhancer (SE)-regulated transcription factors (TF) in human arterial tissues. **(B)** Ranking of the 1,071 arterial SEs. The ranking number and signaling intensity are averaged from the 9 analyzed samples. SEs annotated to transcription factors are highlighted in red and PRDM16 is labeled. **(C)** IGV tracks of DNase-seq, ATAC-seq, H3K27ac and H3K4me1 ChIP-seq, typical enhancers (TEs) and SEs identified in aortic tissues, at human *PRDM16* gene locus. **(D-E)** *PRDM16* expression across different human **(D)** and mouse tissues **(E)** as revealed by GTEx portal and bulk RNA-Seq analysis, respectively. Su.: subcutaneous; Vi.: visceral; Th.: thoracic; Co.: coronary; Ti.: tibial; Ce.: cerebellum; Cor.: cortex; Hi.: Hippocampus; AP: atrial appendage; LV: left ventricle; Me.: medulla; Ske.: skeletal. TPM: transcript per million; FPKM: fragments per kilobase of exon per million mapped fragments. **(F-G)** *Prdm16* expression across different mouse tissues by qRT-PCR **(F)** and Western blotting **(G)** analysis. BAT: brown adipocyte tissue. **(H-I)** UMAP of cell cluster, expression of *PRDM16* and SMC marker *MYH11* revealed by an integrative scRNA-seq dataset of human **(H)**, and mouse aortic tissues **(I)** generated by multiple independent studies. Mod: modulated SMCs; Fibro: fibroblast; EC: endothelial cell; PC: pericyte; MΦ: macrophage; NK: natural killer cell; Pla: plasma cell.

A closer examination of the human *PRDM16* gene locus revealed an enrichment of artery DNase I hypersensitivity signals and ATAC-seq peaks. In combination of analyzing H3K27ac and H3K4me ChIP datasets, we found *PRDM16* gene locus is enriched with typical enhancers (TEs) and 3 large SEs identified by ROSE ^3^ in human arterial tissues (**Figure 1C**). These observations suggested that the chromatin landscape of human *PRDM16* gene locus is highly accessible in arterial tissues. Consistently, data from GTEx portal database showed that human *PRDM16* expression is highly enriched in all 3 arterial tissues including thoracic, coronary and tibial artery. *PRDM16* is lowly expressed in the heart, although in which the function of PRDM16 has been previously reported ^27^ (**Figure 1D**). Similarly, analysis of bulk RNA-Seq of different mouse tissues revealed that *Prdm16* is most abundantly expressed in aortic tissues (**Figure 1E**). This aortic tissue-enriched expression pattern of *Prdm16* was further corroborated by qRT-PCR (**Figure 1F**) and Western blotting analysis (**Figure 1G**) that includes the brown adipocyte tissue (BAT), a tissue in which PRDM16 function was best documented ^24-26^. These results together indicate that PRDM16 expression is enriched in arterial tissues in both human and mouse.

To further define the cell types that express *PRDM16* in blood vessels, we examined its expression in the integrative scRNA-seq datasets of human and mouse aortic tissues, respectively. Both datasets recapitulated major vascular cell types such as normal SMCs (SMC), phenotypically modulated SMC clusters (Mod), fibroblast (Fibro), endothelial cells (EC), and immune cells including T cells, B cells, macrophages (MΦ) and natural killer (NK) cells (**Figure 1H-I & Online Figure S1-2**). This in silico analysis revealed that *PRDM16* is predominantly expressed in normal contractile SMCs and some ECs, and down-regulated in phenotypically modulated SMCs, in aortic tissues of both human and mouse (**Figure 1H-I**). To further clarify the expression of PRDM16 in heart, we re-analyzed publicly available scRNA-seq data generated in embryonic and adult heart of both human (**Online Figure S3**) and mouse (**Online Figure S4**). These analyses confirmed the *PRDM16* expression is enriched in SMCs, in addition to ECs, cardiomyocytes and adipocytes in heart of both human and mouse. Re-analysis of scRNA-seq of SMC-enriched intestinal tissues from both human and mouse revealed that *PRDM16* expression is absent in visceral SMCs (**Online Figure S5**). Taken together, these findings suggest that the CAD gene PRDM16 is preferentially expressed in arterial tissues and VSMCs but not in visceral SMCs, suggesting a previously undocumented function of PRDM16 in VSMCs.

### Loss of *Prdm16* in SMCs alters transcriptomes of VSMCs in mouse aorta

As an initial step to explore the VSMC function of PRDM16 in vivo, we deleted *Prdm16* specifically in SMCs in adult mice by breeding *Prdm16*^*F/F*^ mice ^24^ with mice expressing inducible Cre recombinase under control of SMC-specific *Myh11* promoter ^29^ (**Figure 2A**). This strategy allows for generating *Prdm16* null alleles in SMCs due to a frameshift mutation caused by excision of floxed exon 9 that is mediated by tamoxifen-induced Cre activation. Adult mice were injected with tamoxifen to initiate Cre activation, followed by 2 weeks of washout period, and aortic tissues were harvested for histological and bulk RNA-Seq analysis after additional 60 days (**Figure 2B**). The specific and efficient deletion of *Prdm16* exon 9 in the arteries of *Prdm16* iSM KO mice was validated by detection of the deleted allele with PCR genotyping and was further confirmed by qRT-PCR analysis (**Online Figure S6A-C**). Mice lacking *Prdm16* in SMCs exhibit no obvious gross abnormalities and have comparable body weight to control mice (**Online Figure 6D**). HE staining further demonstrated that *Prdm16* deficiency in SMCs has no discernible effect on the morphology of aortic tissues as evidenced by indistinguishable aortic medial thickness (**Figure 2C-D**) and lumen area compared to control mice (**Online Figure 6E**).

**Figure 2.**
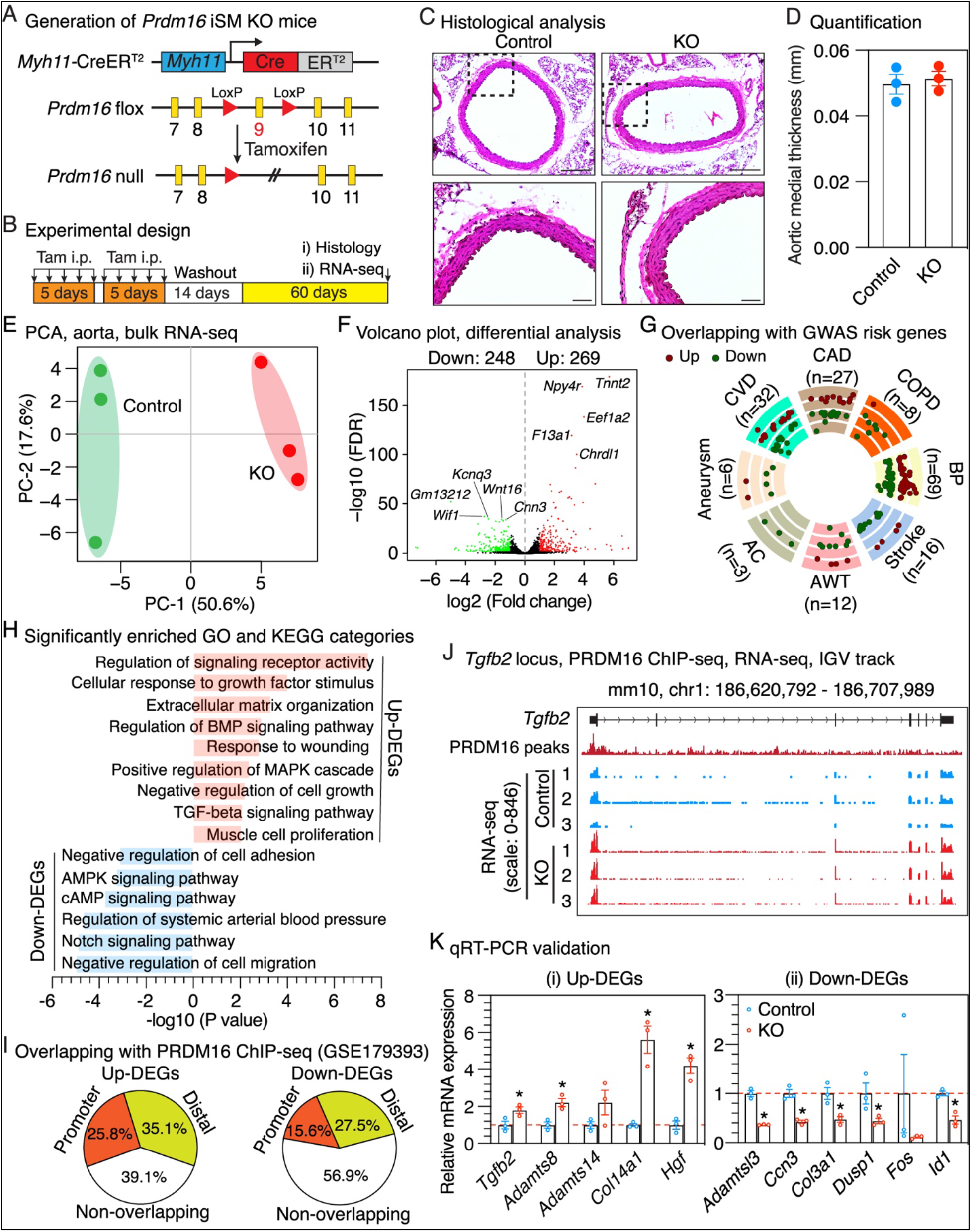
Deletion of *Prdm16* in SMCs in mice alters transcriptome of VSMCs. **(A)** Strategy for generating *Prdm16* inducible smooth muscle-specific KO (iSM KO) mice. **(B)** Schematic diagram of experimental design. Tam: Tamoxifen; i.p.: intraperitoneal injection. **(C)** Representative pictures of HE staining on thoracic aorta from control and *Prdm16* iSM KO mice. The boxed areas are magnified beneath. Scale bar: 50 μm. **(D)** Quantification of the aortic medial thickness. N=3. **(E)** Principal component analysis (PCA) using all the expressed genes identified by bulk RNA-Seq. **(F)** Volcano plot showing the differentially expressed genes (fold change >2 and FDR <0.05) identified by bulk RNA-Seq in aortic tissues of *Prdm16* iSM KO mice as compared to control mice. The top 5 most significantly changed up- and down-regulated genes are labeled. **(G)** Overlapping dysregulated genes identified by bulk RNA-Seq following *Prdm16* deletion with GWAS risk genes for various cardiovascular diseases. CAD: coronary artery disease; COPD: chronic occlusive pulmonary disease; BP: blood pressure; AWT: arterial wall thickness; AC: arterial calcification; CVD: cardiovascular disease. **(H)** Selected enriched GO terms and KEGG pathways for up- and down-regulated genes identified by bulk RNA-Seq following *Prdm16* deletion. **(I)** Common PRDM16-dependent genes identified by bulk RNA-Seq in *Prdm16* iSM KO mouse aorta and a public PRDM16 ChIP-seq data generated in mouse heart. **(J)** IGV tracks of PRDM16 ChIP-seq and bulk RNA-Seq of *Prdm16* iKO vs control aorta showing that the *Tgfb2* gene contains PRDM16 ChIP-seq peaks within its promoter region and is up-regulated in aortic tissues of *Prdm16* iSM KO mice compared to control mice. **(K)** qRT-PCR validation of selected differentially expressed genes identified by RNA-Seq. N=3. *P <0.05; unpaired Student *t* test.

We next sought to assess the effect of *Prdm16* loss on the gene expression programs of VSMCs. We performed bulk RNA-Seq on aortic tissues from both *Prdm16* iSM KO and control mice. As anticipated, examination of RNA-Seq reads demonstrated that the number of reads derived from the floxed exon 9 are dramatically reduced in *Prdm16* iSM KO mouse aorta as compared to that of control mice (**Online Figure 6F**). PCA analysis using all the detected expressed genes revealed KO samples were segregated from control samples (**Figure 2E**), suggesting ablation of *Prdm16* altered the global transcriptome profile of VSMCs. A total of 269 and 248 up- and down-regulated genes were identified following *Prdm16* deletion by differential analysis (Fold change >2 and FDR <0.05) (**Figure 2F and Online Table S4**). Interestingly, 43 of these dysregulated genes have been identified as risk genes by GWAS for various cardiovascular diseases such as CAD, chronic occlusive pulmonary disease (COPD), BP and aneurysm (**Figure 2G and Online Table S4**). Subsequent functional enrichment analysis revealed differentially expressed genes were significantly overrepresented in a wide spectrum of functional categories with some of them are strongly implicated in VSMC phenotypic switching and vascular disease development. For instance, up-regulated genes were involved in response to external stimuli, extracellular matrix organization, MAPK signaling pathway and muscle cell proliferation. Notably, genes involved in BMP and TGF-β signaling pathways were up-regulated in *Prdm16* KO VSMCs (**Figure 2H**). Down-regulated genes are mainly associated with negative regulation of cell adhesion and cell migration, as well as AMPK, cAMP and Notch signaling pathways (**Figure 2H**).

To examine whether these dysregulated genes are direct PRDM16 targets or not, we integrated a PRDM16 ChIP-seq data generated in embryonic mouse heart ^27^. This analysis showed that 60.9% and 43.1% of the up- and down-regulated genes contain PRDM16 ChIP-seq peaks, with 25.8% and 15.6% of them being occupied by PRDM16 within the proximal promoter regions (**Figure 2I and Online Table S4**). Interestingly, we found that *Tgfb2* gene, an upstream effector of TGF-β signaling pathway, is up-regulated upon *Prdm16* depletion and contains PRDM16 ChIP-seq peaks in its promoter region, suggesting *Tgfb2* is likely a direct target of PRDM16 and is repressed by PRDM16 (**Figure 2J**). Furthermore, we selected a list of up-/down-regulated differentially expressed genes including *Tgfb2* and matrix genes (*Adamts8, Adamts14, Adamtsl3, Col14a1* and *Col3a1*), as well as several other genes with little known function in SMCs and vascular diseases such as *Hgf, Fos* and *Id1* for qRT-PCR validation (**Online Figure 6G**). These selected genes are either risk genes for cardiovascular diseases (**Online Figure 6H**) or/and contain PRDM16 ChIP-seq peaks within their genomic region (**Online Figure 6I**). The differential expression of most of the selected genes following *Prdm16* KO was confirmed by qRT-PCR analysis (**Figure 2K**). Collectively, our results suggest that deletion of *Prdm16* in SMCs alters the transcriptome of VSMCs in mice. Specifically, PRDM16 likely negatively regulates TGF-β signaling pathway by inhibiting *Tgfb2* expression through directly binding to the promoter region of *Tgfb2* gene.

## DISCUSSION

In this study, we demonstrate that *PRDM16*, a GWAS identified CAD risk gene with previously well-documented roles in brown adipocytes ^24-26^ and cardiomyocytes ^27^, is enriched with artery SEs and preferentially expressed in arterial tissues and VSMCs, but not in visceral SMCs. Deletion of *Prdm16* gene in SMCs in mice does not affect the aortic morphology but significantly alters expression of many CAD risk genes and genes involved in VSMC phenotypic modulation in aortic tissues, suggesting it is a novel regulator of VSMC homeostasis.

PRDM16 is best known as a master molecular switch of brown fat differentiation ^24-26^ and is recently documented to play a critical role in regulating compaction of cardiomyocytes during murine embryonic development ^27^. In this study, we demonstrated that PRDM16 gene contains numerous artery enhancers and SEs, and is most abundantly expressed in arterial tissues, as compared to other tissues including the brown adipocyte tissue and heart (**Figure 1A-G**). Among vascular cell types, *PRDM16* expression is highly enriched in SMCs as revealed by scRNA-seq of human and mouse aortic tissues (**Figure 1H-I**). This finding is supported by recent scATAC-seq (single-cell sequencing assay for transposase-accessible chromatin) studies of human aortic tissues showing that the *PRDM16* gene locus is enriched with SMC-specific scATAC-seq peaks ^30,31^. PRDM16 is also expressed in ECs with a much lower abundance as compared to SMCs, in agreement with the recent study reporting that PRDM16 plays a role in regulating arterial flow recovery via endothelial function ^28^. Interestingly, unlike other SMC-enriched genes that are expressed in both vascular and visceral SMCs of GI tissues such as MYOCD ^32^, LMOD1 ^33^ and *CARMN* ^34^, PRDM16 exhibits negligible expression in GI tissues and visceral SMCs (**Figure 1D-G & Online Figure S3-5**). Previous studies showed that global or SMC-specific knockout of these pan SMC-enriched genes in mice often leads to lethal GI phenotypes ^35-37^, which largely impedes the efforts for investigating their vascular function. Therefore, the VSMC-enriched expression pattern of PRDM16 offers a unique opportunity to study its function in VSMCs and vascular disorders by inherently circumventing GI phenotypes. The regulatory mechanism conferring the VSMC-enriched expression of PRDM16 is unclear. We speculate it lies in chromatin landscape, as the gene locus contains many chromatin regions that are specifically accessible in VSMCs 30,31. Future studies may focus on unveiling which regulatory element(s) and upstream transcription factor(s) direct the enriched expression of PRDM16 in VSMCs.

We observed that PRDM16 expression is reduced in phenotypically modulated SMCs compared to contractile SMCs in atherosclerosis (**Figure 1H-I**). To test the consequence of loss of PRDM16 in SMCs in vivo, we ablated *Prdm16* specifically in SMCs in mice. Our results show that *Prdm16* deficiency in adult SMCs does not affect the aortic morphology (**Figure 2C-D**), but significantly perturbs the transcriptomic homoeostasis of aortic tissues (**Figure 2E-F**). The dysregulated genes upon *Prdm16* ablation include a subset of known CAD risk genes (**Figure 2G**), suggesting *Prdm16* is not only a CAD gene but also regulates other cardiovascular diseases-associated genes. Furthermore, *Prdm16* deficiency activates expression of genes involved in SMC phenotypic switching, such as those related to response to stimuli, matrix organization and MAPK signaling pathway that is a major regulator of SMC proliferation and migration ^38^ (**Figure 2H**). Meanwhile, genes negatively regulating cell adhesion and migration, as well as Notch signaling pathway that mediates VSMC differentiation and contraction ^39^ are down-regulated following *Prdm16* loss. These findings together implicate that PRDM16 plays a dual role in both promoting differentiation genes and inhibiting de-differentiation genes in VSMCs, reminiscent of its bifunctional role in brown adipocytes and cardiomyocytes where it acts as both transcriptional activator and repressor ^16,27^. The *Prdm16* loss-induced transcriptomic changes likely disrupt VSMC homeostasis and predispose SMCs to vascular diseases such as atherosclerosis and aneurysm.

Among the activated genes following *Prdm16* inactivation is *Tgfb2. Tgfb2* encodes an upstream ligand (TGFB2, transforming growth factor beta-2) of TGF-β signaling pathway and is a GWAS identified risk gene for multiple vascular disease such as aortic aneurysm 40,41, BP ^42^, and COPD ^43^. TGF-β is a well-known pleiotropic regulator of SMC phenotype and function, and either deficient or excess TGF-β signaling may disrupt VSMC homeostasis ^44^. On one hand, TGF-β promotes SMC contraction via canonical SMAD signaling and therefore inactivation of TGF-β pathway impairs SMC contraction and worsens vascular disease development ^40,41,45^. On the other hand, TGF-β increase SMC proliferation, migration and matrix remodeling, thus promoting progression of aortic aneurysm ^46^ and neointimal formation ^47^. Enhanced TGF-β signaling is observed in the media of aortic tissues from aneurysm patients ^48,49^ and animal models ^50^. Consistently, a recent single-nuclear RNA-seq study showed that *TGFB2* expression is activated in SMC clusters of aneurysmal human aorta as compared to that of healthy control ^51^. We reasoned that TGFB2 is a direct and bona fide target of PRDM16 because 1) it harbors PRDM16 ChIP-seq peaks within its proximal promoter region (**Figure 2J**); 2) increased *Tgfb2* expression is consistently observed following *Prdm16* deletion in cardiomyocytes in multiple independent studies ^27,52,53^, a similar finding to our study. Thus, all these observations together pointed that *Tgfb2* is the primary target underlying the function of *Prdm16* in regulating SMC homeostasis. Furthermore, because TGFBs are secretary cytokines and well-known regulators of pro-fibrosis/EndMT (endothelial-to-mesenchymal transition) ^54,55^ and immune cell function ^56^, TGFB2 activation induced by *Prdm16* loss may have paracrine effects on other vascular cell types, in addition to its intrinsic function in VSMCs.

In summary, our study demonstrates that the CAD gene *PRDM16* is preferentially expressed in aortic tissues and VSMCs and is a novel regulator of VSMC homeostasis. Together with previous study showing multiple CAD SNPs residing in SMCs-specific peaks at *PRDM16* gene locus and are associated with *PRDM16* expression ^31^, we speculate that PRDM16 contributes to CAD progression mainly through regulating VSMC phenotype, although non-SMCs such as brown fat adipocytes and endothelial cells can be additional potential contributor to disease development ^57-60^. Future studies are warranted to explore its function under pathological conditions by introducing vascular diseases, such as atherosclerosis, to *Prdm16* iSM KO mice.

## ONLINE SUPPLEMENTAL DATA

Online methods Online Table S1-4 Online Figure S1-6

## AUTHOR CONTRIBUTIONS

K. Dong and J. Zhou conceptualized the project and designed the experiments. K. Dong, G. Hu and Y. Yao performed the experiments. K. Dong, X. He and J. Zhou performed the analysis of the data. K. Dong and J. Zhou wrote the original draft and the final version of the manuscript. All authors read and approved the final paper.

