## Supplementary material for "Coronary Artery Disease Risk Gene *PRDM16* is Preferentially Expressed in Vascular Smooth Muscle Cells and a Potential Novel Regulator of Smooth Muscle Homeostasis": Online Methods, Table S1-3 and Figure S1-6

**Kunzhe Dong, PhD**

Immunology Center of Georgia

Department of Pharmacology & Toxicology

Medical College of Georgia

Augusta University

1462 Laney Walker Blvd.,

Augusta GA 30907

**Jiliang Zhou, MD/PhD**

Department of Pharmacology & Toxicology

Medical College of Georgia

Augusta University

CB-3628 (Office); CB-3606 (Lab)

1459 Laney Walker Blvd

Augusta, GA 30912

### Online Methods

#### Identification of human artery SEs

The aligned Bam and narrowPeak files of H3K27ac ChIP-seq data generated from 9 different human arterial tissues were downloaded from ENCODE database (**Online Table S1**) and used to identify artery SEs by using ROSE program [1] with the default arguments. Super-imposed SEs that are present in at least 5 artery samples were selected and further ranked based on a combination of average ranking and intensity across the 9 samples. For each SE, the associated genes were annotated using R package ChIPseeker [2] and intersected with 1,839 human transcription factors obtained from SCENIC database [3].

#### Bioinformatics analysis of public bulk RNA-Seq and scRNA-seq data

Bulk RNA-Seq data of aortic tissue [4] and 10 other tissues [5] of mouse were used to evaluate *Prdm16* abundance indicated by FPKM (fragments per kilobase of exon per million mapped fragments) across different mouse tissues.

To examine *PRDM16* expression at the single-cell level, we generated a merged scRNA-seq dataset of aortic tissues by integrating multiple public scRNA-seq datasets from independent studies for both human (including GSE131778 [6], GSE155512 [7] and GSE155468 [8]) and mouse (including GSE174384 [9], GSE117963 [10], GSE131776 [6] and GSE155513 [7]) using the R package Harmony which has been shown to reduce technical batch effects [11]. Briefly, the indicated datasets were SCTransformed and merged using the merge command of Seurat package [12]. The RunHarmony command was then used to integrate these datasets. UMAP coordinates, neighbors and clusters were then calculated with the reduction parameter set to ‘harmony’. FindAllMarkers function in Seurat was used to identify marker genes for each cluster.

scRNA-seq data of embryonic (Study # SCP1021) and adult (Study # SCP498) human heart [13] including the UMAP coordinate, cell type, gene expression of each cell was obtained from BROAD Institute Single Cell Portal. UAMP visualization of cell cluster and gene expression was generated by custom R scripts. Count matrix of scRNA-seq for embryonic (E10.5) [14] and non-cardiac cells from adult mouse heart [15] were downloaded from GEO (GSE122403) and ArrayExpress database (E-MTAB-6173), respectively, and processed using Seurat package as we previously described [16]. The processed scRNA-seq data of adult mouse heart was obtained from Tabula Muris [17] and visualized using Seurat package [12]. scRNA-seq datasets generated in embryonic human gut (https://www.gutcellatlas.org/) [18] and adult human colon (GSE156905) [19], as well as mouse colon and ileum (Single Cell Portal, SCP1038) were re-analyzed as we previously described [20].

**Quantitative reverse transcription-PCR (qRT-PCR) analysis**

Total RNA from mouse tissues was isolated with TRIzol reagent (Invitrogen). 0.8 μg of RNA was used as template for reverse transcription (RT) with random hexamer primers using the High Capacity RNA-to-cDNA kit (Invitrogen). Real time PCR was performed with SYBR green PCR master mix (Applied Biosystems) and respective primers listed in **Online Table S2**. All samples were amplified in duplicates. Relative gene expression was converted using the $2^{-\triangle\triangle CT}$ method against the internal control house-keeping gene glyceraldehyde-3-phosphate dehydrogenase (*Gapdh*) where $\triangle\triangle CT = (CT_{experimental\ gene} - CT_{experimental\ Gapdh}) - (CT_{control\ gene} - CT_{control\ Gapdh})$.

**Protein extraction and Western blotting**

Protein lysates were extracted from different mouse tissues by RIPA buffer (Thermo Fisher Scientific) plus 1% protease/phosphatase inhibitor cocktail (Thermo Fisher Scientific). After sonication and centrifugation of the tissue lysates, proteins in the supernatant were quantified by BCA assay (Thermo Fisher Scientific) and resolved on a 7.5% SDS-PAGE gel with 15 μg per lane where appropriate. Primary antibodies against PRDM16 (R&D, AF6295, sheep, 1:1000), ACTA2 (Sigma, A2547, mouse, 1:5000), and GAPDH (Santa Cruz Biotechnology, sc-32233, mouse, 1:1000) were used. Secondary antibodies conjugated with horseradish peroxidase were then used to visualize the target proteins in blot. Image was acquired by AMERSHAM ImageQuant 800 biomolecular imager.

**Mouse breeding and genotyping**

The *Prdm16* flox mice with exon 9 flanked with loxP sites [21] were purchased from the Jackson Laboratory (Stock #: 024992). *Myh11*-CreER$^{T2}$ transgenic mice (originally generated by Dr. Stefan Offermanns's lab as previously published [22]) were obtained from Dr. Joseph Miano at the University of Rochester (now at Augusta University). All strains have been maintained on a C57BL/6 background. To generate *Prdm16* inducible SM-specific KO (iSM KO) mice, female mice homozygous for *Prdm16* flox allele (*Prdm16*$^{F/F}$) were crossed with male *Myh11*-CreER$^{T2}$ mice. Subsequently the *Myh11*-CreER$^{T2+}$; *Prdm16*$^{F/W}$ male mice were bred with *Prdm16*$^{F/W}$ female mice to generate *Myh11*-CreER$^{T2+}$; *Prdm16*$^{F/F}$ mice and *Myh11*-CreER$^{T2+}$; *Prdm16*$^{W/W}$ mice that serve as control. At 8 weeks of age, mice from both groups were intraperitoneally injected with tamoxifen (1 mg/mouse/day) for 10 days with 2 days' break between the first and second 5 injections, followed by a washout of tamoxifen for 2 weeks. Mice were monitored for additional 60 days daily for signs of phenotypes and then sacrificed for histological and bulk RNA-seq analysis. Only male mice were used in this study because *Myh11*-CreER$^{T2}$ transgene is located in the Y chromosome [22].

**Sections and Hematoxyline/Eosin (HE) staining**

Mice were euthanized by an overdose of 4% Isoflurane via inhalation, then systemically perfused with PBS via the left ventricle. Isolated thoracic aortic tissues were fixed with 4% paraformaldehyde in PBS overnight at 4°C, washed 3 times with PBS, then kept in 30% sucrose in PBS overnight at 4°C. Fixed tissues were embedded in optimal cutting temperature compound (OCT) and kept at -80°C till cryo-sectioning. Sections were cut at 8 μm thickness and HE staining was performed following standard protocol as previously described [16]. HE-stained images were captured using an ECHO Revolution RVL2-K inverted microscope. Sections were analyzed blindly by an independent investigator for aortic media layer thickness and lumen area using Image J software.

**Transcriptome analysis by bulk RNA-Seq**

Aortic tissues of 3 *Prdm16* iSM KO and 3 control mice were isolated and adventitial layer was removed under a stereoscope. Total RNA from the aortic tissues was then extracted using TRIzol reagent (Invitrogen) and subjected to whole transcriptome RNA-seq analysis at the Genome Technology Access Center at Washington University. Sequencing libraries were constructed from purified RNA using RiboErase kit (Kapa Biosystems) according to the manufacturer's instructions. Libraries were sequenced on a NovaSeq 6000 system (Illumina) using paired-end reads extending 150-bp bases.

Obtained RNA-seq reads were then mapped and quantitated to Ensembl release 101 mouse reference genome (mm10) with an Illumina DRAGEN Bio-IT. Raw count for each gene was rounded and the integer count was used for subsequent analysis. Only genes with count >10 in all the samples of at least one group were considered as expressed genes and used for subsequent analysis. Differential expression analysis was performed with R package DEseq2 [23]. Cutoff values of fold change greater than 2 and false discovery rate (FDR) less than 0.05 were considered statistically significant between control and KO groups. Principle component analysis (PCA) and volcano plot was generated using custom R script. GO (Gene Ontology) and KEGG pathway analysis was carried out by Metascape (http://metascape.org).

**Integrative analysis of GWAS and bulk RNA-Seq**

GWAS identified risk genes for different cardiovascular diseases including CAD, chronic obstructive pulmonary disease (COPD), blood pressure (BP), stroke, arterial wall thickness (AWT), artery calcification (AC), aneurysm and cardiovascular disease (CVD) were obtained from GWAS Catalog database (https://www.ebi.ac.uk/gwas/). Differentially expressed genes in mouse aorta following *Prdm16* deletion were intersected with the obtained gene list to identify *Prdm16*-regualed risk genes for different cardiovascular diseases. The overlapping results was visualized with R package GOplot [24].

**PRDM16 ChIP-seq data de novo analysis**

PRDM16 ChIP-seq peaks generated in C57BL/6 mouse heart were obtained from GEO database (GSE179371) [25]. The peaks were annotated using ChIPseeker package [2] and overlapped with differentially expressed genes identified in the aortic tissues of *Prdm16* iSM KO mice. Raw reads of PRDM16 ChIP-seq for whole heart tissues of WT mice were downloaded and aligned against Ensembl mouse reference genome mm10 using Bowtie 2 [26]. The generated Bam files were used for Integrative Genomics Viewer (IGV) visualization.

**Statistical analysis**

GraphPad Prism (version 9.2.0) was used for the statistical analysis. All data are expressed as mean ± SEM of at least 3 independent experiments. Tests used for statistical significance evaluations are specified in figure legends. An unpaired 2-tailed *t* test was used for data involving 2 groups only. Values of $P < 0.05$ were considered statistically significant for qRT-PCR analysis and FDR-adjusted $P < 0.05$ was used as the threshold for statistical significance for bulk RNA-seq analysis.

**Online Table S1**
**Information of H3K27ac ChIP-seq data of human arterial tissues in ENCODE used for identification of artery super-enhancers.**

| *Tissue* | *Sample ID* | *ChIP-seq group* | *ENCODE ID* | |
|---|---|---|---|---|
| | | | **Bam file** | **narrowPeak file** |
| Aorta | Aorta 1 | H3K27ac | ENCFF265HDL | ENCFF072EQH |
| | | Control | ENCFF861FUK | |
| | Aorta 2 | H3K27ac | ENCFF434DCE | ENCFF064PQH |
| | | Control | ENCFF278NFU | |
| Ascending aorta | As. Aorta 1 | H3K27ac | ENCFF823SXL | ENCFF020COG |
| | | Control | ENCFF893ZJC | |
| | As. Aorta 2 | H3K27ac | ENCFF128VGR | ENCFF208DZK |
| | | Control | ENCFF299AOC | |
| Tibial artery | T. Artery 1 | H3K27ac | ENCFF140QKE | ENCFF134QRV |
| | | Control | ENCFF259JSG | |
| | T. Artery 2.1 | H3K27ac | ENCFF437QEP | ENCFF121HDP |
| | T. Artery 2.2 | H3K27ac | ENCFF459ELZ | |
| | | Control | ENCFF400VJM | |
| Thoracic aorta | Th. Aorta 1 | H3K27ac | ENCFF946NGS | ENCFF595RQJ |
| | | Control | ENCFF901HZC | |
| | Th. Aorta 2 | H3K27ac | ENCFF816GNU | ENCFF121HDP |
| | | Control | ENCFF924RQL | |

**Online Table S2**

**List of primers used for qRT-PCR (F: forward; R: reverse).**

| *Gene name* | *Primer name* | *Sequence (5'-3')* | *Application* |
|---|---|---|---|
| *Prdm16* | P1 (F) | TAGTGTGTAGCTGCTTCTGGGCTCA | qRT-PCR, detection of exon 1/2 |
| | P2 (R) | ACAGGATGCCGTCTTCGGTCTCCT | |
| | P9 (F) | TGCCTAAGGTGTGCCCAGCACAGC | qRT-PCR, detection of exon 9/10 |
| | P10 (R) | CGCAGGTACTTCTCTTTCAGGACTC | |
| *Tgfb2* | F | TTGTTGCCCTCCTACAGACTGG | qRT-PCR |
| | R | GTAAAGAGGGCGAAGGCAGCAA | |
| *Adamts8* | F | TCATGCAACACAGAGGAATGTCCAC | qRT-PCR |
| | R | CTCTGCAAAACAGCTTGCATCGGTC | |
| *Adamts14* | F | TTCCACAGGTTCCACTGGTCTCGCT | qRT-PCR |
| | R | GCATTGCTCATCCATGGAGTAGTCG | |
| *Col14a1* | F | GAGGTTCAACTTCAGGCTTGTGCGC | qRT-PCR |
| | R | GGCATTCAAGTGCCACTCTATTCTG | |
| *Hgf* | F | GACGGTATCCATCACTAAGAGTGGC | qRT-PCR |
| | R | CTTCTTCCCCTCGAGGATTTCGACA | |
| *Adamtsl3* | F | TTTGCGGAGATGTCTGACTGG | qRT-PCR |
| | R | ACTGCACGTCATTGTAGGCTG | |
| *Ccn3* | F | AGATGAGACCCTGTGACCAGAGCAG | qRT-PCR |
| | R | GCAGAAGTACTGACAGTTCGGCTCA | |
| *Col3a1* | F | CCTGGCTCAAATGGCTCAC | qRT-PCR |
| | R | CAGGACTGCCGTTATTCCCG | |
| *Dusp1* | F | CTACCAGTACAAGAGCATCCCTGTG | qRT-PCR |
| | R | CTCATGAGGTAAGCAAGGCAGATGG | |
| *Fos* | F | CGGGTTTCAACGCCGACTA | qRT-PCR |
| | R | TGGCACTAGAGACGGACAGAT | |
| *Id1* | F | CTGCTCTACGACATGAACGGCTGCT | qRT-PCR |
| | R | TCAGCGACACAAGATGCGATCGTCG | |
| *Gapdh* | F | GGCATTGCTCTCAATGACAA | Internal control for qRT-PCR analysis |
| | R | TGTGAGGGAGATGCTCAGTG | |

Notes: location of primer P1, P2, P9 and P10 for *Prdm16* gene is illustrated in Online Figure S6A. All the primers are designed for mouse genes.

**Online Table S3**

**List of primers used for genotyping (F: forward; R: reverse).**

| *Primer name* | *Sequence (5'-3')* |
|---|---|
| F | CATGGTTCACATGGTCAAGACCAC |
| R1 | CACAGTCCTTGCACTTGATCTGCGT |
| R2 | AGAGCTGCAGGGAGATTGACAAGTG |

Notes: location of primer F, R1 and R2 was illustrated in Online Figure S6A.

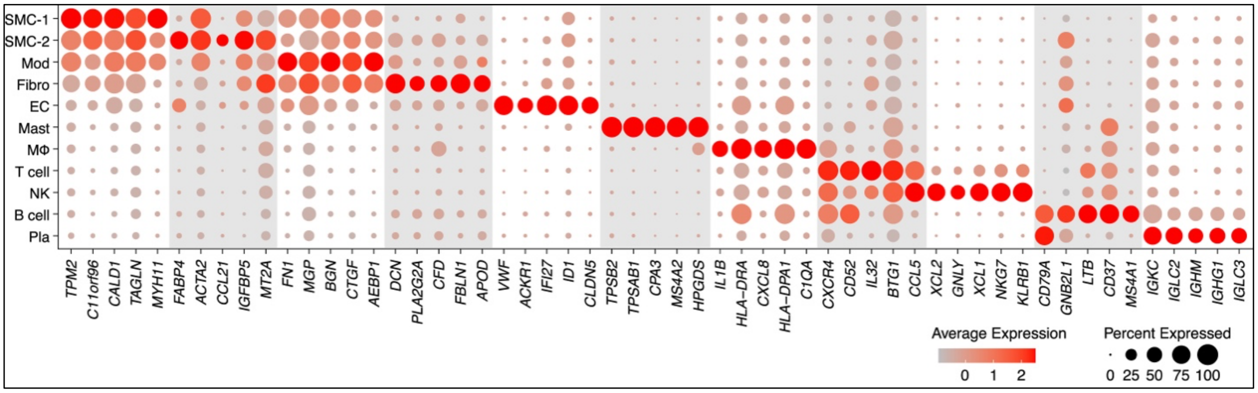


**Online Figure S1. Dot plot showing the top 5 genes defining each cell cluster for the merged scRNA-seq data of human arterial tissues (GSE131778, GSE155512, GSE155468).** Mod: modulated SMCs; Fibro: fibroblast; EC: endothelial cell; MΦ: macrophage; NK: natural killer cell; Pla: plasma cell.

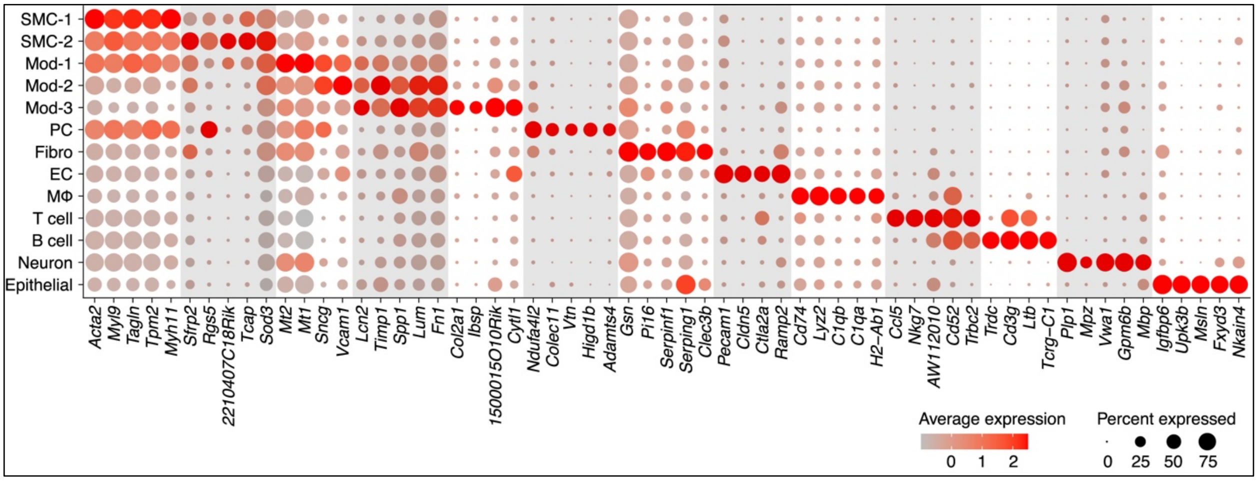


**Online Figure S2. Dot plot showing the top 5 genes defining each cell cluster for the merged scRNA-seq datasets of arterial tissues of normal and atherosclerotic mouse models (GSE174384, GSE117963, GSE131776, GSE155513).** Mod: modulated SMCs; PC: pericyte; Fibro: fibroblast; EC: endothelial cell; MΦ: macrophage.

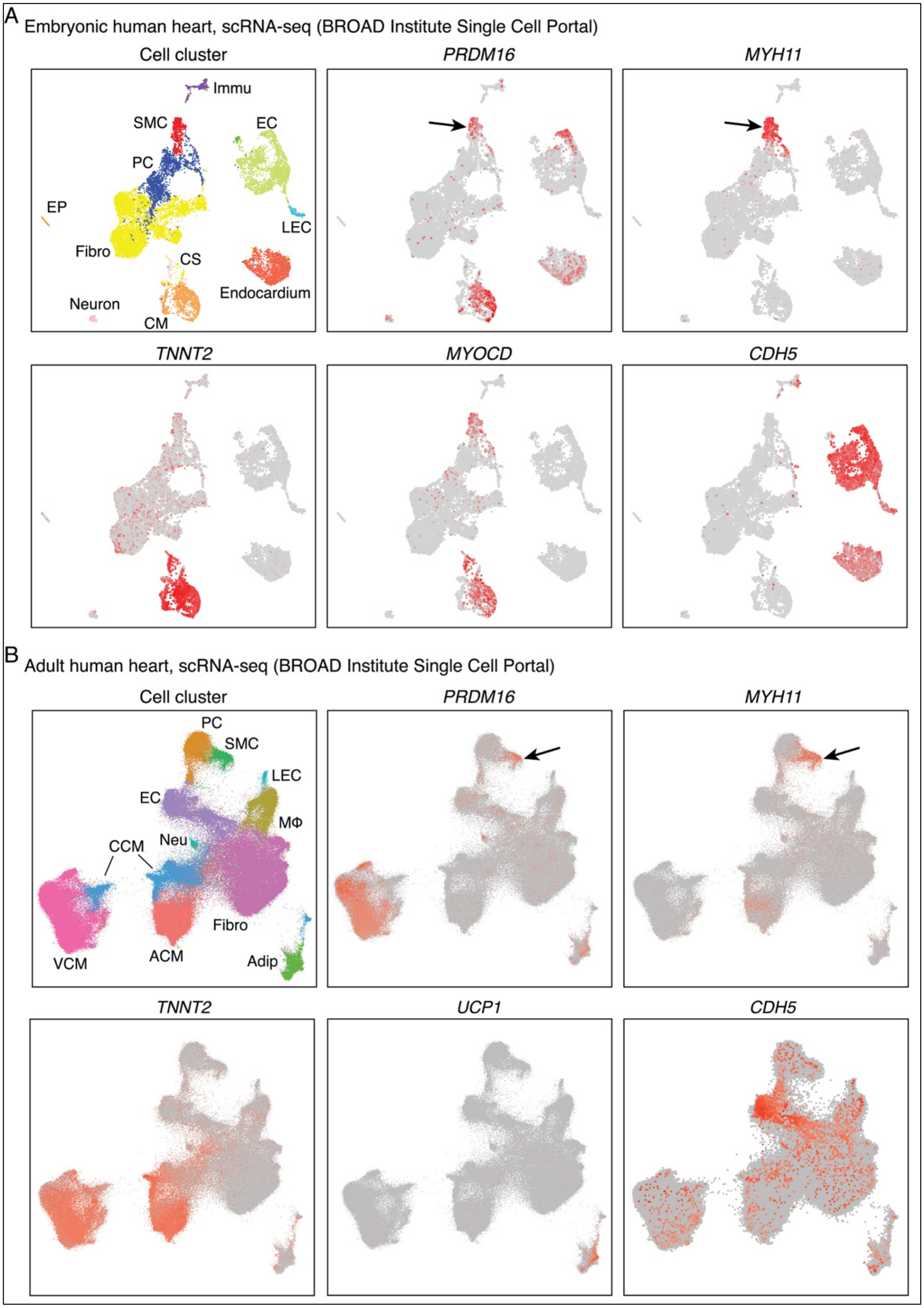
A
Embryonic human heart, scRNA-seq (BROAD Institute Single Cell Portal)
Cell cluster
PRDM16
MYH11
Immu
SMC
EC
PC
EP
LEC
Fibro
CS
Endocardium
Neuron
CM
TNNT2
MYOCD
CDH5
B
Adult human heart, scRNA-seq (BROAD Institute Single Cell Portal)
Cell cluster
PRDM16
MYH11
PC
SMC
LEC
EC
MΦ
Neu
CCM
Fibro
VCM
ACM
Adip
TNNT2
UCP1
CDH5

**Online Figure S3. PRDM16 expression in embryonic (A) and adult human heart (B) as revealed by scRNA-seq analysis.** EC: endothelial cell; LEC: lymphatic endothelial cell; PC: pericyte; CS: conduction system; CM: cardiomyocyte; EP: epicardium; Immu: immune cells; Fibro: fibroblast; MΦ: macrophage; VCM: ventricular cardiomyocyte; ACM: atrial cardiomyocyte; CCM: cytoplasmic cardiomyocyte; Neu: neuronal cell; Adip: adipocyte. *MYH11*, *TNNT2*, *CDH5* and *UCP1* are used as markers for SMCs, CMs, ECs and adipocytes, respectively. *MYOCD* is used as markers for both SMCs and CMs. Both datasets were downloaded from BROAD Institute Single Cell Portal (links are provided in the Methods) and original annotation of cell types are used. Arrows point to SMC cluster.

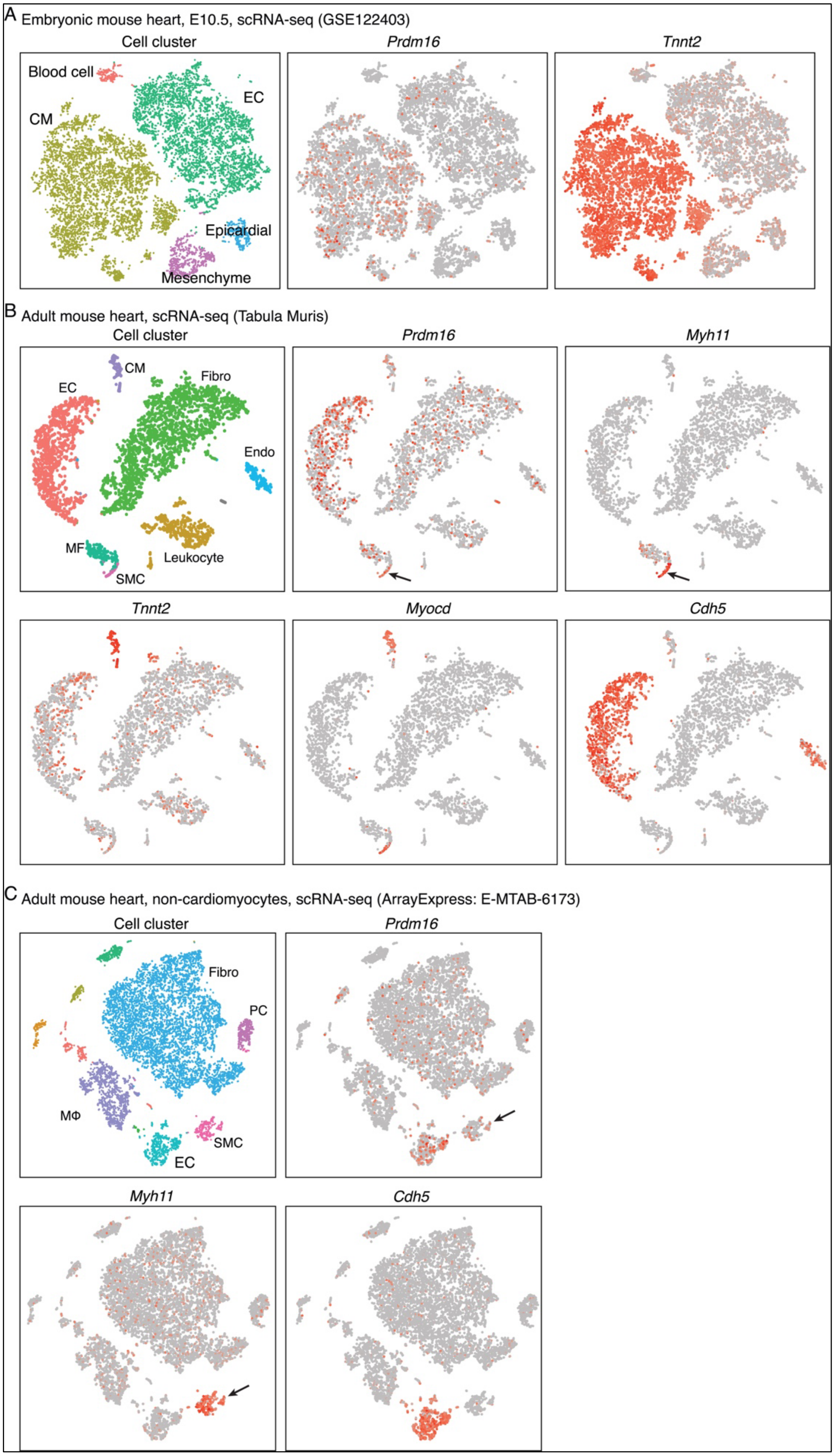
A Embryonic mouse heart, E10.5, scRNA-seq (GSE122403)
Cell cluster
Prdm16
Tnnt2
Blood cell
EC
CM
Epicardial
Mesenchyme
B Adult mouse heart, scRNA-seq (Tabula Muris)
Cell cluster
Prdm16
Myh11
CM
Fibro
EC
Endo
MF
Leukocyte
SMC
Tnnt2
Myocd
Cdh5
C Adult mouse heart, non-cardiomyocytes, scRNA-seq (ArrayExpress: E-MTAB-6173)
Cell cluster
Prdm16
Fibro
PC
MΦ
SMC
EC
Myh11
Cdh5

**Online Figure S4. *Prmd16* expression in (A) embryonic and (B-C) adult mouse heart as revealed by scRNA-seq analysis.** EC: endothelial cell; CM: cardiomyocyte; Fibro: fibroblast; MΦ: macrophage; MF: myofibroblast cell; Endo: endocardial cell; PC: pericyte. *Myh11*, *Tnnt2* and *Cdh5* were used as markers for SMCs, CMs and ECs, respectively. *Myocd* was used as markers for both SMCs and CMs. The original annotation for cell types in adult heart from Tabula Muris database was used. Arrows point to SMC cluster.

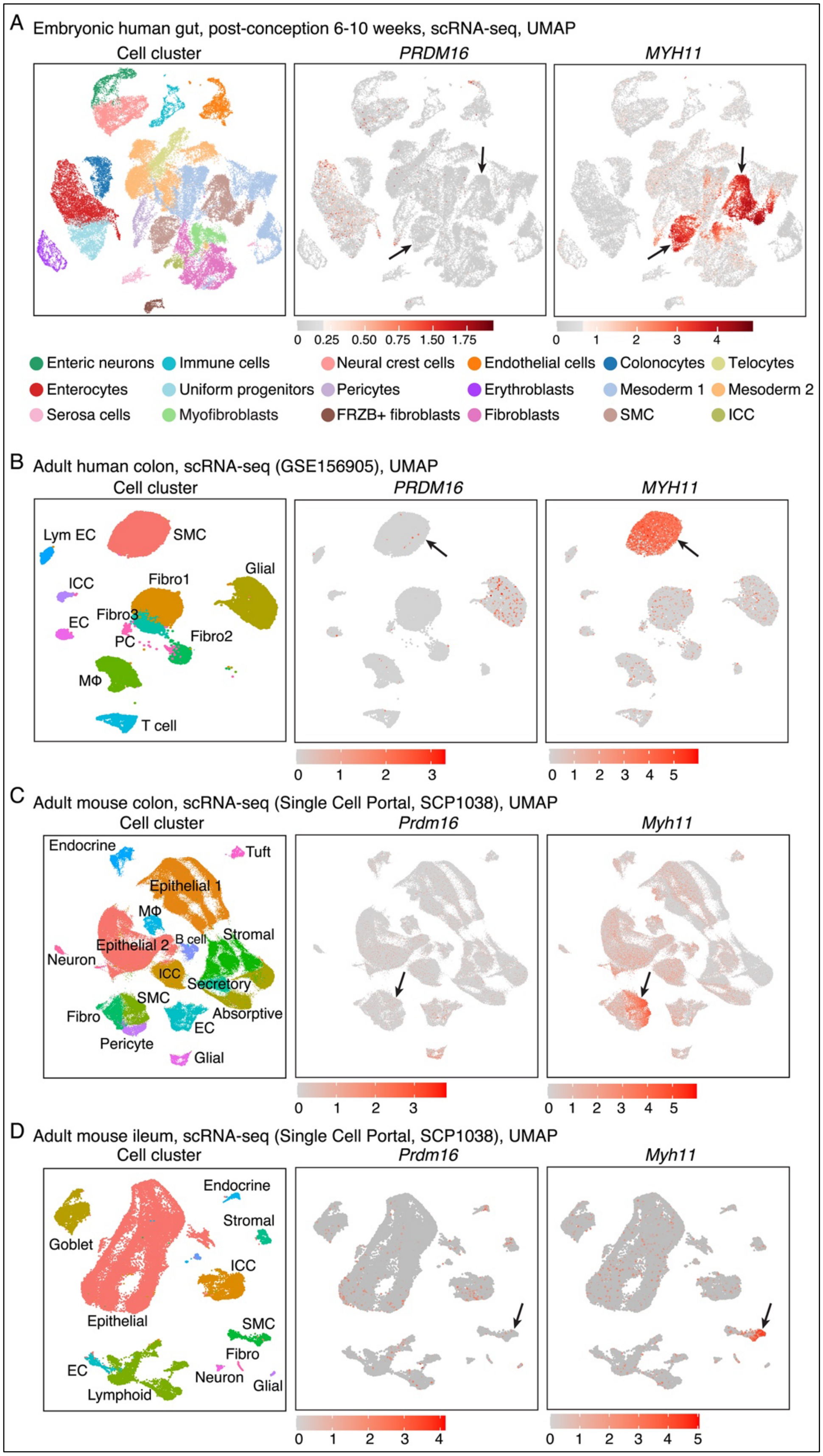

A Embryonic human gut, post-conception 6-10 weeks, scRNA-seq, UMAP
Cell cluster
PRDM16
MYH11
0 0.25 0.50 0.75 1.50 1.75
0 1 2 3 4
Enteric neurons
Immune cells
Neural crest cells
Endothelial cells
Colonocytes
Telocytes
Enterocytes
Uniform progenitors
Pericytes
Erythroblasts
Mesoderm 1
Mesoderm 2
Serosa cells
Myofibroblasts
FRZB+ fibroblasts
Fibroblasts
SMC
ICC
B Adult human colon, scRNA-seq (GSE156905), UMAP
Cell cluster
PRDM16
MYH11
Lym EC
SMC
Glial
ICC
Fibro1
EC
Fibro3
PC
Fibro2
MΦ
T cell
0 1 2 3
0 1 2 3 4 5
C Adult mouse colon, scRNA-seq (Single Cell Portal, SCP1038), UMAP
Cell cluster
Prdm16
Myh11
Endocrine
Tuft
Epithelial 1
MΦ
Epithelial 2
B cell
Stromal
Neuron
ICC
Secretory
SMC
Fibro
Absorptive
EC
Pericyte
Glial
0 1 2 3
0 1 2 3 4 5
D Adult mouse ileum, scRNA-seq (Single Cell Portal, SCP1038), UMAP
Cell cluster
Prdm16
Myh11
Endocrine
Stromal
Goblet
ICC
Epithelial
SMC
Fibro
EC
Neuron
Glial
Lymphoid
0 1 2 3 4
0 1 2 3 4 5

**Online Figure S5. *PRDM16* expression in human and mouse gastrointestinal tissues as revealed by scRNA-seq. (A)** *PRDM16* expression revealed by scRNA-seq of embryonic human gut (post-conception 6-10 weeks), **(B)** adult human colon, **(C)** adult mouse colon and **(D)** ileum tissues. ICC: interstitial cells of Cajal; PC: pericyte; Lym EC: Lymphatic endothelial cell; Fibro: Fibroblast; MΦ: Macrophage. Arrows point to SMC clusters.

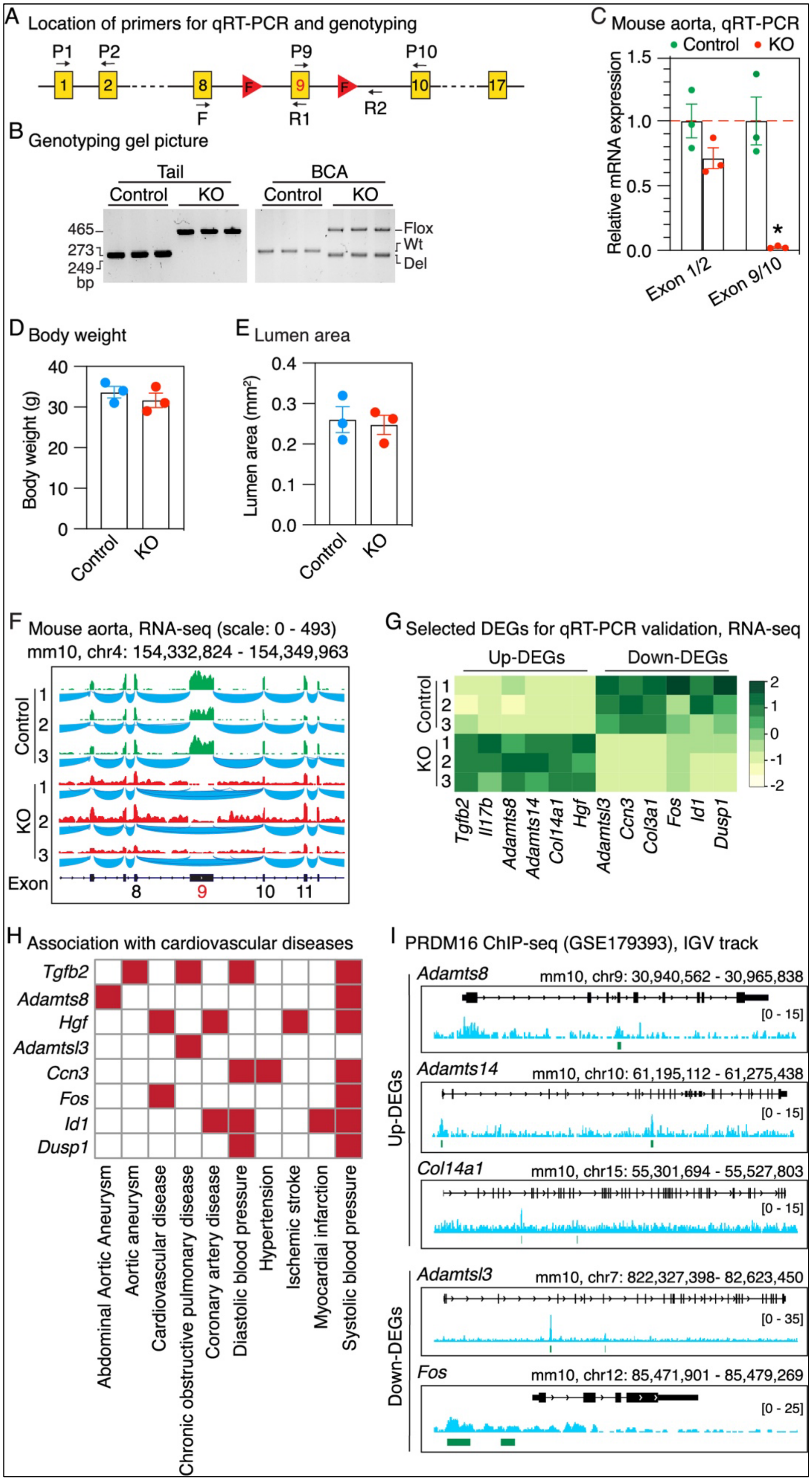
A Location of primers for qRT-PCR and genotyping
P1 P2 P9 P10
1 2 8 F 9 F 10 17
F R1 R2
B Genotyping gel picture
Tail BCA
Control KO Control KO
465 273 249 bp
Flox Wt Del
C Mouse aorta, qRT-PCR
Control KO
Relative mRNA expression
Exon 1/2 Exon 9/10
D Body weight
Body weight (g)
Control KO
E Lumen area
Lumen area (mm²)
Control KO
F Mouse aorta, RNA-seq (scale: 0 - 493)
mm10, chr4: 154,332,824 - 154,349,963
Control KO Exon 8 9 10 11
G Selected DEGs for qRT-PCR validation, RNA-seq
Up-DEGs Down-DEGs
Control KO
Tgfb2 Il17b Adamts8 Adamts14 Col14a1 Hgf Adamtsl3 Ccn3 Col3a1 Fos Id1 Dusp1
H Association with cardiovascular diseases
Tgfb2 Adamts8 Hgf Adamtsl3 Ccn3 Fos Id1 Dusp1
Abdominal Aortic Aneurysm
Aortic aneurysm
Cardiovascular disease
Chronic obstructive pulmonary disease
Coronary artery disease
Diastolic blood pressure
Hypertension
Ischemic stroke
Myocardial infarction
Systolic blood pressure
I PRDM16 ChIP-seq (GSE179393), IGV track
Up-DEGs Down-DEGs
Adamts8 mm10, chr9: 30,940,562 - 30,965,838 [0 - 15]
Adamts14 mm10, chr10: 61,195,112 - 61,275,438 [0 - 15]
Col14a1 mm10, chr15: 55,301,694 - 55,527,803 [0 - 15]
Adamtsl3 mm10, chr7: 822,327,398- 82,623,450 [0 - 35]
Fos mm10, chr12: 85,471,901 - 85,479,269 [0 - 25]

**Online Figure S6. Histological and bulk RNA-Seq analysis of aortic tissues of *Prdm16* inducible SMC-specific KO mice as compared to control mice. (A)** Schematic diagram illustrating the location of primers used for *Prdm16* gene qRT-PCR (P1/P2 for detecting exon 1 and 2, P9 and P10 for detecting exon 9 and 10) and genotyping (F, R1 and R2). **(B)** Representative agarose gel picture of PCR genotyping using DNA extracted from tail and brachiocephalic artery (BCA) as the template. **(C)** qRT-PCR analysis of *Prdm16* expression in aortic tissues of *Prdm16* iSM KO and control mice, using primers across exon 1 and 2, as well as exon 9 and 10 (PCR amplicon is depicted in "**A**"), respectively. N=3; *P<0.05; Unpaired student *t* test. **(D)** Body weight of *Prdm16* iSM KO mice and control mice prior to sacrifice. N=3. **(E)** Quantification of lumen area for aortic sections of *Prdm16* iSM KO mice and control mice. N=3. **(F)** Distribution of RNA-Seq junction reads surrounding *Prdm16* exon 9 showing dramatic reduction in number of junction reads spanning *Prdm16* exon 8-9, and exon 9-10 in *Prdm16* iSM KO mice as compared to control mice. **(G)** Heatmap showing the up- and down-DEGs that were identified by RNA-Seq and selected for qRT-PCR validation. **(H)** Heatmap showing association of the indicated genes with cardiovascular diseases identified by GWAS studies. **(I)** Integrative Genomics Viewer (IGV) visualization of PRDM16 occupancy at selected target gene loci. Green bars indicate the significant PRDM16 ChIP-seq peaks identified by the original study.
